# Enabling supratheoretical isopropanol yields from carbon-negative glucose fermentations with *Clostridium acetobutylicum*-*Clostridium ljungdahlii* cocultures

**DOI:** 10.1101/2025.07.14.664808

**Authors:** Noah B. Willis, Jonathan K. Otten, Hyeongmin Seo, Pradeep C. Munasinghe, John D. Hill, Eleftherios T. Papoutsakis

## Abstract

Synthetic microbial cocultures, which combine the unique capabilities of multiple microbes into one process, have significant potential for sustainable production of fuels and chemicals. Most studies of defined cocultures have tested relatively low cell densities in lab-scale batch cultures, not the high cell density fed-batch or continuous processes with cell retention typically required to achieve industrially-relevant volumetric productivities. Here, we explore the impact of increased cell density on isopropanol production from the syntrophic coculture of genetically-modified *Clostridium acetobutylicum* [CACas9 Δ*hbd* (p95ace02_atoB), with deleted 4-C metabolism expressing an acetone-formation pathway on the plasmid] with WT *Clostridium ljungdahlii* using first a pseudo-perfusion approach followed by perfusion culture. CACas9 Δ*hbd* (p95ace02_atoB) produces acetone without any 4-C metabolites and *C. ljungdahlii* converts that acetone to isopropanol. To explore the mechanism by which these cultures enable supratheoretical isopropanol yields, we first identified NADH-driven hydrogen conversion in CACas9 Δ*hbd* (p95ace02_atoB) as the thermodynamically-limiting step for acetone and thus isopropanol production. We then demonstrated the ability of *C. ljungdahlii* to mitigate this issue by eliminating detectable hydrogen accumulation in the coculture. Pseudo-perfusion cocultures showed that high cell densities combined with a high population fraction of *C. ljungdahlii* enable dramatic increases in isopropanol yields beyond the thermodynamic limitation imposed in CACas9 Δ*hbd* (p95ace02_atoB) monocultures. Finally, we demonstrate carbon-negative fermentation of glucose to isopropanol as the sole alcohol product in a perfusion bioreactor.

## 1. INTRODUCTION

There is significant interest in the use of synthetic, defined microbial cocultures for the sustainable production of fuels and chemicals. Industrial-scale microbiological production processes have typically relied on monocultures of well-characterized model organisms (often with significant genetic modifications) or undefined consortia of wild-type microbes (such as in wastewater treatment and food manufacturing). Little work has been done with cocultures using fed-batch or perfusion growth modes, which are typically required to achieve the high cell densities, and correspondingly high volumetric productivities, necessary for economic viability.

Since cocultures inherently rely on optimized interspecies interactions, and interspecies cell distance decreases (on average) with increasing cell density, it is logical to assume that a given microbial consortium may perform much differently at high cell densities than at low cell densities, making both absolute culture cell density and relative species abundance crucial operational parameters. Formation of biofilms, a ubiquitous and ancient method to increase local cell density, has been shown to be highly relevant for the formation, function, and resilience of natural and infectious microbial communities [1]. There is increasing evidence the last few years that in some cases, synthetic cocultures can perform direct exchange of cytoplasmic material via nanotube-like structures or interspecies cell-to-cell contact (reviewed in [2]). Our group has extensively shown that interspecies contact in the coculture pairing of *Clostridium acetobutylicum* and *Clostridium ljungdahlii* leads to large scale exchange of protein, RNA, and DNA and the generation of interspecies hybrid cells [3–5]. The impact of varied absolute and relative cell densities on the unique behavior of our *C. acetobutylicum-C. ljungdahlii* pairing specifically, and on the many other interesting behaviors observed in microbial cocultures generally, remains largely unexplored. One would assume that absolute cell densities and relative species abundance are crucial tuning parameters for maximizing these types of syntrophic interactions.

In addition to interspecies exchange of cellular material and cell fusion, the *C. acetobutylicum-C. ljungdahlii* coculture also presents several other useful phenotypes of practical significance. Specifically, this coculture achieves improved carbon recovery and alcohol yield per mol of consumed glucose relative to either monoculture [3]. The *C. acetobutylicum-C. ljungdahlii* pairing also synthesizes significant quantities of isopropanol, a valuable chemical that neither coculture partner produces independently. *C. acetobutylicum* produces acetone which *C. ljungdahlii* converts to isopropanol. Importantly, separating the two species by a permeable membrane significantly impairs isopropanol formation and the carbon recovery of glucose into metabolites [3]. These findings suggest that interspecies proximity and cell-to-cell contact are crucial for isopropanol production and perhaps other useful phenotypes. These results also suggest that maximizing interspecies proximity and cell-to-cell contact, via optimizing the species ratio and culturing the consortia in a high-density reactor system, may strengthen the syntrophic interactions between *C. acetobutylicum* and *C. ljungdahlii*, potentially producing further increases in the fraction of substrate carbon into metabolites and, notably, isopropanol.

In this study, in order to increase acetone production (and thus isopropanol production in coculture), we used an engineered *C. acetobutylicum* strain, CACas9 Δ*hbd* [6]. The strain was further modified by transforming it with plasmids that carry two versions of a synthetic acetone-formation pathway. Using these strains in monocultures and cocultures with *C. ljungdahlii*, we studied the impact of high density coculture fermentation and species ratio on isopropanol yields from glucose (mol isopropanol formed per mol glucose used). First, we identify a thermodynamic constraint impacting this isopropanol yield that makes it impossible to produce acetone as the sole product from glucose using *C. acetobutylicum* monocultures without an engineered electron sink. Next, we identify coculture with an acetogen, such as *C. ljungdahlii*, as a strategy to overcome this constraint based on *C. ljungdahlii’s* ability to “remove” electrons from *C. acetobutylicum*. We then show that high density, “pseudo-perfusion” co-cultures, specifically those with high *C. ljungdahlii* population fractions, can further improve yields to enable supratheoretical isopropanol yields from glucose beyond the maximum biological yield in *C. acetobutylicum* monocultures. Finally, we demonstrate that these supratheoretical yields can be maintained stably for over 100 hours in a perfusion bioreactor, enabling carbon-negative glucose fermentation to isopropanol as the sole alcohol product with strong volumetric productivity.

## 2. MATERIALS AND METHODS

### Plasmid and *C. acetobutylicum* strain construction

The strains and associated plasmids used in this study are listed in Table 1. The list of primers and gBlocks used to construct these strains are described in Table S1. The p95ace02_atoB plasmid designed and used in this study was constructed using NEBuilder HiFi DNA assembly (New England Biolabs, MA, USA). The plasmid backbone was amplified without the *C. acetobutylicum thl* gene using p95ace02a as a template. The *atoB* insert was ordered as a gBlock fragment (Integrated DNA Technologies, IA, USA) codon optimized for *C. acetobutylicum* with assembly overhangs compatible with the amplified plasmid backbone. The PCR reaction to amplify the backbone was performed with Phusion DNA polymerase or AccuPrime DNA polymerase (Thermo Fisher Scientific, MA, USA). Plasmids were sequence confirmed via colony PCR using Phire DNA polymerase (Thermo Fisher Scientific, MA, USA) and Sanger sequencing. Transformation of *C. acetobutylicum* was performed as previously described [6].

**Table 1.** List of bacterial strains and plasmids used in this study.

| Name | Descriptions | Source |
| --- | --- | --- |
| **Plasmids** |  |  |
| pAN3 | Km<sup>R</sup>; $\Phi 3T$ I methylase | Al-Hinai et al. (2012) |
| p95ace02a | pta promoter from *C. ljungdahlii* upstream of *ctfA* and *ctfB*; P<sub>synTHL</sub> promoter upstream of *thl* and *adc*; ColE1 ori; *repL*; Amp<sup>R</sup>; MLS<sup>R</sup> | Seo et al. (2024) |
| p95ace02_atoB | Replaces *C. acetobutylicum* *thl* on p95ace02a with *E. coli* *atoB* | This study |
| p100ptaHALO | pta promoter from *C. ljungdahlii* upstream of *Halotag* fluorescent reporter; ColE1 ori; *repB*; Amp<sup>R</sup>; MLS<sup>R</sup> | Charubin et al. (2020) |
| ***E. coli* strains** |  |  |
| DH5α | *fhuA2*Δ(*argF-lacZ*)U169<br>*phoA glnV44 80*Δ(*lacZ*)M15<br>*gyrA96 recA1 relA1 endA1*<br>*thi-1 hsdR17* | NEB |
| ER2275 | *hsdR mcrA recA1 endA1* | NEB |
| ***C. acetobutylicum* strains** |  |  |
| CACas9 Δ*hbd* | CACas9 with markerless *hbd* deletion | Seo et al. (2024) |
| ***C. ljungdahlii* strain** |  |  |
| DSM13528 | Wild-type | DSM |

### Coculture preparation for batch and resuspension fermentations

For *C. acetobutylicum* CACas9 *Δhbd* p95ace02_atoB, a frozen stock was streaked onto a 2xYTG agar plate supplemented with 30mM of sodium butyrate and 40 μg/ml clarithromycin and left at 37°C in an anaerobic incubator for 3 days to allow for colony growth and spore formation. All subsequent culturing steps were performed in Turbo CGM [3] growth medium supplemented with 30 mM sodium butyrate (“Turbo CGMB”) with 100 μg/ml clarithromycin. After incubation, a single colony was inoculated into a test tube containing 10 mL of Turbo CGMB growth medium, heat shocked at 80°C to kill non-sporulated cells and grown statically in an anaerobic incubator at 37°C for 16-24 hours. The resulting pre-culture was used to inoculate 30 mL of Turbo CGMB growth medium to an OD_600_ of 0.1. This culture grew statically in the anaerobic incubator at 37°C until it reached exponential phase (OD_600_ between 0.5-4.0). This pre-culture was used to inoculate the co-culture.

For *C. ljungdahlii* p100ptaHALO, a frozen stock was inoculated into 20 mL of YTF medium [7] supplemented with 3.5 g/L arginine and 25 g/L MES (“YTAF-MES”) with 100 μg/ml clarithromycin in a 160 mL serum bottle which had been pre-flushed for 2 min with an 80% H_2_, 20% CO_2_ gas mix. The serum bottle was pressurized to 20 psig with the 80/20 gas mixture and grown for 24 hours at 37°C on a rotating platform at 90 rpm. This pre-culture was used to inoculate pre-flushed serum bottles each containing Turbo CGMB with 40 grams per liter of glucose and 100 μg/ml clarithromycin to an OD_600_ of 0.05-0.10. These pre-cultures were grown to exponential phase (OD_600_ of 0.6-1.2) and used to inoculate the cocultures.

For coculture preparation, volumes of exponential *C. acetobutylicum* and *C. ljungdahlii* pre-cultures corresponding to the desired target starting optical density and an R value of 5 (ratio of *C. ljungdahlii* to *C. acetobutylicum* cells based on OD_600_) [3] was centrifuged at 5000 rpm, 4°C for 10 min and then resuspended in 20 mL of Turbo CGMB in pre-flushed serum bottles. For batch serum bottle experiments, the target starting optical density was 1.0 OD_600_ and the initial serum bottle pressure was ∼8 psig. For the resuspension serum bottle experiments, the target starting optical density was 6.0 OD_600_ and the initial serum bottle pressure was ∼20 psig.

### Metabolite and coculture population ratio analyses

High pressure liquid chromatography (HPLC) was used to quantify supernatant sugar and solvent concentrations as previously reported [8–10]. For headspace analysis, gas samples were quantified using an Agilent 7890A gas chromatograph equipped with a Supelco 60:80 Carboxen-1000 column and TCD according to the column manufacturer’s recommended protocol for H_2_ and CO_2_. Coculture population fractions of *C. acetobutylicum* and *C. ljungdahlii* were determined using RNA-FISH as previously described [11].

### Coculture perfusion fermentation

Perfusion cocultures were conducted in a bioreactor system consisting of a 3 L glass vessel, bioreactor control tower (Applikon BioConsole ADI1025), and packed bed column (constructed in-house) filled with Pro-Pak 0.16” distillation packing connected in series. 1.2 L working volume was maintained in the reactor vessel and 2.4 L working volume was maintained in the packed column for a total working volume of 3.6 L. An anaerobic environment was maintained via sparging 80/20 H_2_/CO_2_ mixture at 3.6 liters per hour through the packed bed. Reactor pH was controlled via controlled addition of 4M NaOH. To initiate the perfusion coculture, 80 mL of *C. acetobutylicum* pre-culture was inoculated into the perfusion reactor initially containing Turbo CGMB with 60 g/L of glucose and 100 ug/ml clarithromycin. After inoculation, perfusion was initiated; Turbo CGMB with 20 g/L of glucose was fed at 4.86 mL/min and removed at the same rate (approximately 2 vessel volumes per day). Cell retention was performed with a GE Healthcare hollow-fiber cartridge with 0.1 μm pore size and 850 cm^2^ surface area (CFP-1-E-4X2MA). 120 mL of concentrated *C. ljungdahlii* pre-culture was inoculated into the reactor system after 70 hr of fermentation. The reactor was operated in complete cell retention mode until 100 hr, at which point a 10% bleed was initiated.

## 3. RESULTS & DISCUSSION

### 3.1. Knockout of 4-carbon metabolism and overexpression of acetone pathway genes in *C. acetobutylicum* redirect carbon use from butanol to ethanol

As recently reported [6], as a first step to repurpose the *C. acetobutylicum-C. ljungdahlii* coculture for isopropanol production (Fig. 1A), we engineered a *C. acetobutylicum* strain for acetone production by combining two key interventions: knockout of 4-carbon metabolism and overexpression of the acetone pathway genes using strong, constitutive promoters. Briefly, to eliminate butyrate and butanol production (i.e., 4-carbon metabolism), we utilized an engineered *C. acetobutylicum* strain with an integrated inducible Cas9 (for CRISPR editing) to knock out the 3-hydroxybutyryl-CoA dehydrogenase (*hbd*) gene thus generating the “CACas9 Δ*hbd*” strain [6]. Strong and sustained growth of this strain *C. acetobutylicum* strain requires the presence of small concentrations (typically 30 mM) of butyrate as documented [6].

**Figure 1:**
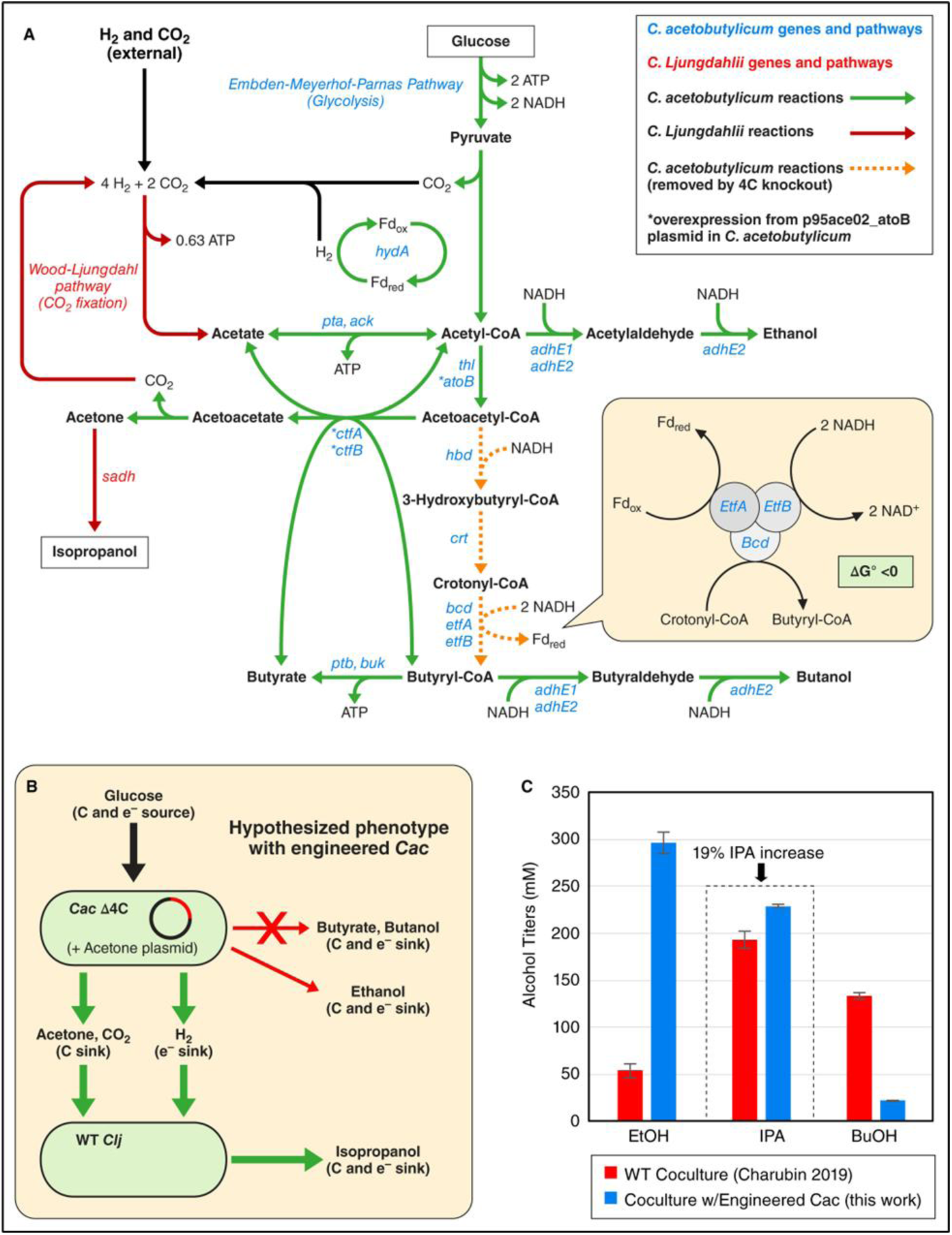
A) Schematic of major metabolic reactions in the coculture system, B) schematic depicting coupled reduction of crotonyl-CoA and ferredoxin by NADH via the Bcd-EtfAB protein complex, C) hypothesized carbon and electron distribution in coculture using 4-carbon knockout strain of *C. acetobutylicum* engineered for acetone production (CACas9 Δ*hbd* p95ace02_atoB), and D) comparison of actual alcohol titers between cocultures using wild-type *C. acetobutylicum* and 4-carbon knockout strain of *C. acetobutylicum* engineered for acetone production (CACas9 Δ*hbd* p95ace02_atoB).

To enhance acetone formation (Fig. 1A), we transformed CACas9 Δ*hbd* with a plasmid, “p95ace02_atoB” (Fig S1A), coding for acetone production plasmid, which is a modified version of the previously reported plasmid p95ace02 [6]. The acetone production pathway is made up of *atoB,* an *Escherichia coli* thiolase; *ctfA/B,* the genes coding for the two chains of the *C. acetobutylicum* Co-A transferase(CoAT); and *adc,* coding for the *C. acetobutylicum* acetoacetate decarboxylase, using two strong promoters, the phosphotransacetylase promoter (P_PTA_) from *C. ljungdahlii* and an engineered *C. acetobutylicum* thiolase promoter (P_synTHL_) [6, 12]. The only difference between p95ace02_atoB and p95ace02 is the use of the *E. coli* thiolase instead of a *C. acetobutylicum* thiolase. We used the *atoB* version of this plasmid (“p95ace02_atoB”) because we have observed improved acetone production in CACas9 Δ*hbd* (p95ace02_atoB) relative to CACas9 Δ*hbd* (p95ace02a) (Fig. S1B) likely because, unlike the *E. coli* thiolase, the *C. acetobutylicum* thiolase is conformationally deactivated by a disulfide bridge under sufficiently oxidized intracellular conditions [13].

As an initial test for isopropanol production, we performed a simple batch coculture between CACas9 Δ*hbd* p95ace02_atoB and *C. ljungdahlii.* For this experiment (and all subsequent coculture experiments reported in this study), *C. ljungdahlii* was transformed with the p100ptaHalo plasmid (expressing the HaloTag fluorescent protein) due to its antibiotic resistance to clarithromycin (required for maintaining the p95ace02_atoB in CACas9 Δ*hbd*). Despite engineering *C. acetobutylicum* to knock out 4-carbon metabolism and overexpress the acetone pathway genes, this co-culture system produced only 19% more isopropanol than our previously demonstrated cocultures [3] of WT *C. acetobutylicum* and *C. ljungdahlii* (Fig. 1D). In the coculture with WT *C. acetobutylicum,* butanol (132 mM, 528 C-mM; C-mM stands for Carbon mM) is a significant carbon sink. In the coculture with the engineered *C. acetobutylicum,* glucose carbon was not redirected towards acetone, as we hypothesized (Fig. 1C), but instead to ethanol (296 mM, 592 C-mM). A small amount of butanol (24 mM) was produced in the engineered coculture, but this was due to conversion of the small amount of butyrate (30 mM) added to the medium to support CACas9 Δ*hbd* growth [6], and, thus, did not derive from glucose carbon.

### 3.2. Constraints on the interconversion of electron carriers limit the acetone yield and explain the increased ethanol production by CACas9 Δ*hbd* (p95ace02_atoB)

Production of large quantities of ethanol by the engineered coculture, despite minimal ethanol production in the coculture with WT *C. acetobutylicum*, is explained by electron balances and thermodynamic constraints on the interconversion between electron carriers in *C. acetobutylicum*. For every mol of glucose catabolized through glycolysis, *C. acetobutylicum* generates two mol of NADH and two mol of reduced ferredoxin (Fd_red_) (Fig. 1A, Table 2). Acetone production does not require any reducing equivalents downstream of the acetyl-CoA node, so, to maximize acetone yields, the reducing equivalents from glucose catabolism to acetyl-CoA are ideally converted to molecular H_2_. In *C. acetobutylicum* monocultures, this H_2_ is released as a waste product. In coculture, electrons from molecular H_2_ are recaptured by *C. ljungdahlii. C. ljungdahlii* uses these electrons to reduce CO_2_ to acetate through the Wood-Ljungdahl Pathway (WLP) (generating energy for growth) and to reduce acetone to isopropanol. Fd_red_, the electron donor for *C. acetobutylicum’s* hydrogenase [14] (Fig. 1A, Table 2), can be readily converted to molecular H_2_. NADH, on the other hand, must first be converted to Fd_red_ for H_2_ generation.

**Table 2.**
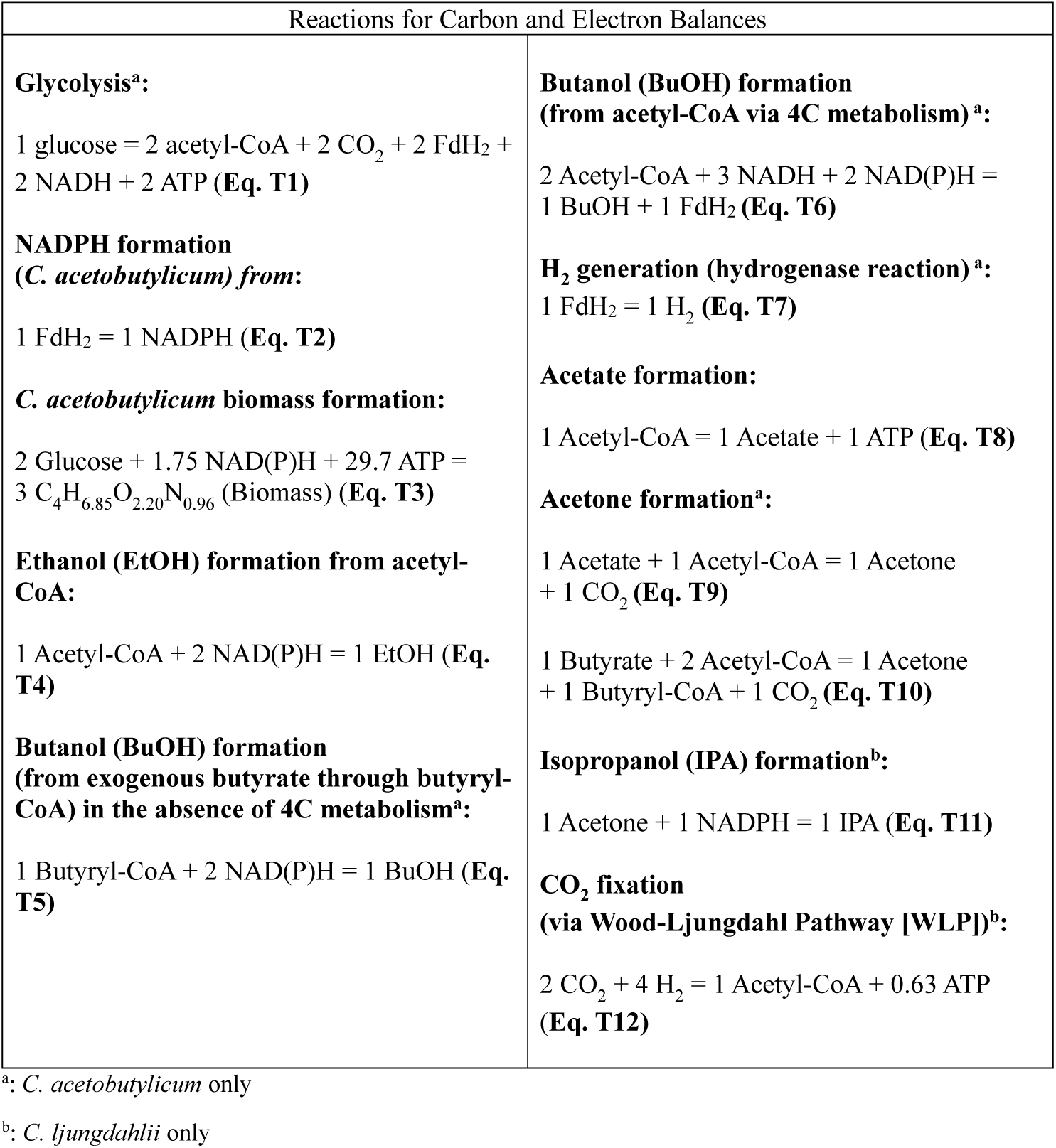
Stoichiometric carbon and electron balances for core metabolic reactions in *C. acetobutylicum* and *C. ljungdahlii*.

The major enzyme complex responsible for Fd_red_:NADH interconversion in *C. acetobutylicum* is the Bcd-EtfAB complex [15] (Fig. 1A). Bcd (butyryl-CoA dehydrogenase) is responsible for crotonyl-CoA reduction to butyryl-CoA. EtfA and Bcd are electron-transport proteins which facilitate the energetic coupling of electron-bifurcating redox reactions [16]. The corresponding three genes are part of the operon which codes for the pathway for converting acetoacetyl-CoA to butyryl-CoA (Fig. 1A). The *C. acetobutylicum* Bcd-EtfAB complex can convert Fd_red_ to NADH in the absence of crotonyl-CoA [15]. However, the reverse reaction (NADH conversion to Fd_red_) is endergonic (ΔG > 0) [17]. One way to overcome this energetic barrier is to couple the endergonic reduction of oxidized ferredoxin (Fd_ox_) to Fd_red_ by NADH with the exergonic reduction – catalyzed by the Bcd-EtfAB complex – of crotonyl-CoA to butyryl-CoA by NADH [18].

In view of these limitations, in *C. acetobutylicum*, NADH can only be converted to Fd_red_ (and then to H_2_) in the presence of crotonyl-CoA. Since our engineered *C. acetobutylicum* strain, CACas9 *Δhbd* p95ace02_atoB, has the 4-carbon metabolism knocked out, it does not synthesize crotonyl-CoA and thus cannot perform the reduction of Fd_ox_ by NADH (and thus to produce H_2_ from NADH). Therefore, production of ethanol is the only available NADH sink in this strain. Since each mol of glucose consumed produces 2 mol of NADH from glycolysis, and each mole of ethanol produced requires 2 mol of NADH (Fig. 1A, Table 2), *at least one mol of ethanol must be produced per mol glucose consumed to satisfy the electron balance when NADH cannot be converted to H_2_*. This is depicted schematically in Figure S2. In monocultures of CACas9 *Δhbd* p95ace02_atoB grown under a nitrogen headspace, we observed 1.02 mol ethanol per mole glucose (in addition to <0.1 mol butanol per glucose from exogenous butyrate reduction) (Fig. 2B), consistent with the predicted minimum yield of ethanol. Since, in addition to the two mol of NADH, each mol of ethanol requires one mole of acetyl-CoA, only one mole of acetyl-CoA per glucose remains available for acetone production, resulting in a maximum yield of 0.5 mol of acetone per mol of glucose (Eq. 1, derived using the equations of Table 1).

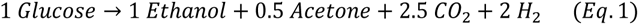

**Figure 2:**
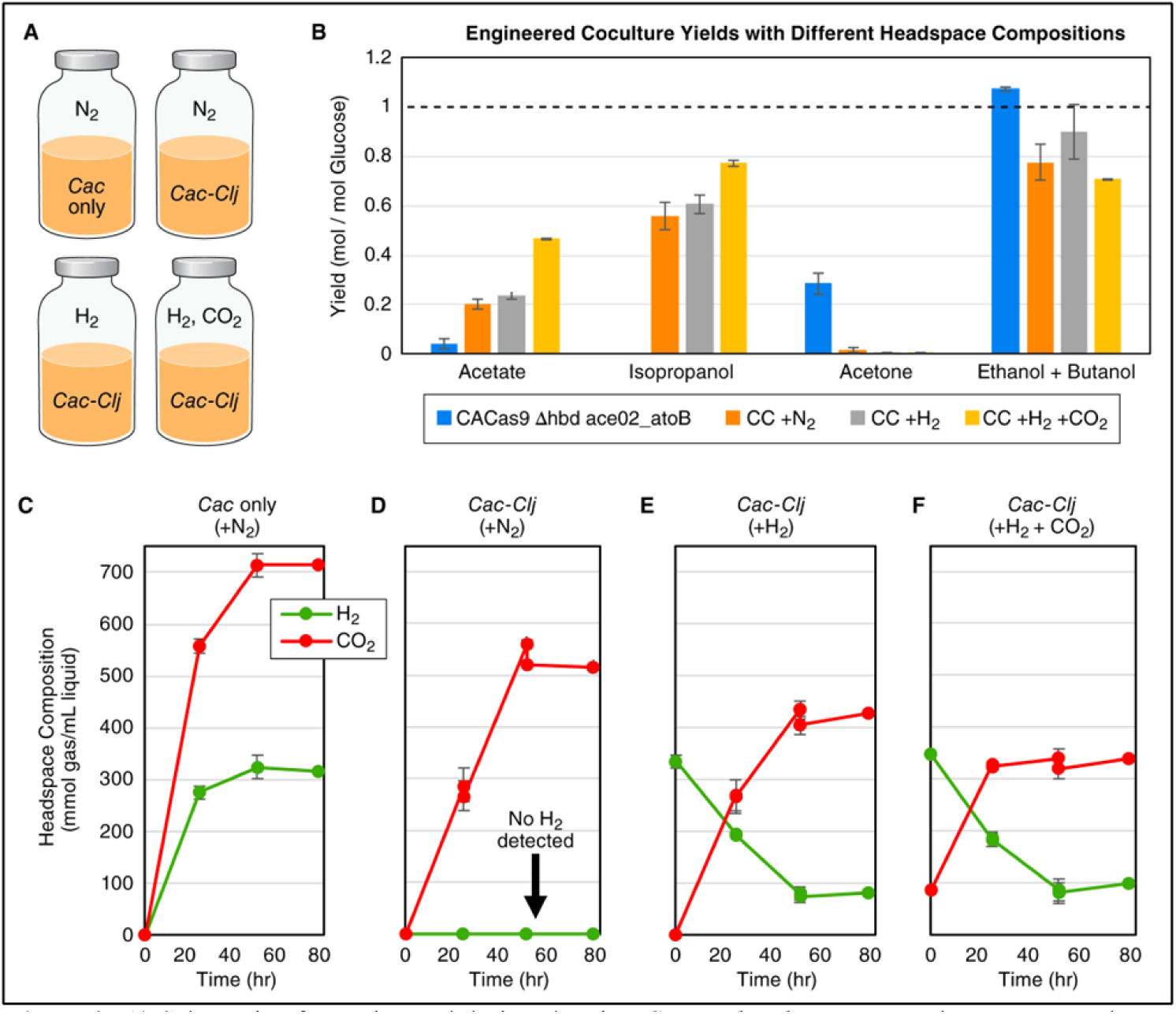
A) Schematic of experimental design showing *C. acetobutylicum* monoculture grown under N2 and *C.acetobutylicum-C. ljungdahlii* cocultures grown under N2, H2, and H2/CO2. B) Per glucose yields of major metabolites for the *C. acetobutylicum* monoculture (under N2 atmosphere) and three coculture conditions (under N2, H2, and H2/CO2). C) headspace gas composition kinetics for *C. acetobutylicum* monoculture grown under N2. D) Headspace gas composition kinetics for *C. acetobutylicum-C. ljungdahlii* coculture grown under a N2 atmosphere. E) Headspace gas composition kinetics for *C. acetobutylicum-C. ljungdahlii* coculture grown under H2. F) Headspace gas composition kinetics for *C. acetobutylicum-C. ljungdahlii* coculture grown under H2/CO2. Error bars represent standard deviations from two biological replicates.

Retaining 4-carbon metabolism (which generates crotonyl-CoA from glucose) enables enough conversion of NADH to H_2_ to increase the maximum acetone yield per glucose by 20% (0.6 mol acetone per mol glucose) (Eq. 2, derived using the equations of Table 2).

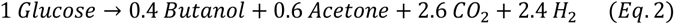

With or without the 4-carbon metabolism, we refer to this requirement – which fundamentally limits acetone yields – to use acetyl-CoA to re-oxidize NADH as a “thermodynamic limitation” because it is caused by the endergonic nature of the reaction that converts NADH to Fd_red_ and eventually to H_2_.

### 3.3. *C. ljungdahlii* eliminates H_2_ accumulation in coculture with *C. acetobutylicum* and extracts additional electrons from exogenous H_2_

Although monocultures of 4-carbon knockout *C. acetobutylicum* strains cannot convert NADH to H_2_, it may be possible to overcome this limitation by coculturing *C. acetobutylicum* with *C. ljungdahlii*. It is well-established that certain organisms which cannot convert NADH to H_2_ when grown in monoculture can be induced to convert NADH to H_2_ in coculture with a partner H_2_-consuming organism, so long as this partner organism has a sufficiently low H_2_ threshold. “H_2_ threshold” refers to the minimum partial pressure at which an organism can utilize H_2_ [19]. An organism with a low H_2_ threshold can utilize H_2_ even at extremely low partial pressures. For example, when grown on glucose, monocultures of *Clostridium thermocellum* produce 2 mol of H_2_ per mol glucose and a mix of ethanol and lactate to oxidize the additional 2 mol of NADH produced via glycolysis per mol glucose utilized. However, when cocultured with *Methanothermobacter thermoautotrophicum*, per mol glucose, *C. thermocellum* generates 4 mol of H_2_, which is quickly consumed by *M. thermoautotrophicum* along with CO_2_ to make methane. [20]. This occurs because methanogens like *M. thermoautotrophicum*, due to their extremely low H_2_ thresholds [21], can reduce the H_2_ partial pressure to extremely low levels. When the H_2_ partial pressure is sufficiently low, NADH to H_2_ conversion becomes exergonic (ΔG < 0) [17]. For this reason, we hypothesized that increasing the cell density and the fraction of *C. ljungdahlii* cells in coculture may enable decreased ethanol production (and corresponding increase in acetone yield) from *C. acetobutylicum* by one (or both) of two mechanisms: by removing electrons (in the form of NADH, FdH_2_ and/or H_2_) from *C. acetobutylicum* directly via cytoplasmic exchange [4], and/or by enabling *C. acetobutylicum* to convert NADH to H_2_ by maintaining low H_2_ partial pressures in the *C. acetobutylicum* microenvironment.

As a first step to explore *C. ljungdahlii’s* ability to remove electrons from *C. acetobutylicum*, we measured the soluble and gaseous metabolite kinetics in cocultures, using our engineered *C. acetobutylicum* strain, with three different starting headspace compositions: pure N_2_, pure H_2_, and an 80% H_2_/20% CO_2_ mixture. We also included a *C. acetobutylicum* monoculture under N_2_ as an control (Fig. 2A). The media for these experiments contained starting sugar concentrations of 80 g/L glucose and 5 g/L fructose. To stimulate the growth of *C. acetobutylicum*, the medium contained also 30 mM of butyrate [6].

For metabolite analyses, we present the summed yields (mol) of ethanol and butanol per mol glucose and refer to this value as the “primary alcohol yield” (to distinguish it from the yield of isopropanol, a secondary alcohol). Reduction of 1 mol acetyl-CoA to 1 mol ethanol requires 2 mol NADH (Table 2). Similarly, reduction of 1 mol butyrate (exogenously added in the case of 4-carbon knockout strains) to 1 mole butanol also requires 2 mol of NADH (Table 2). Therefore, since the production of 1 mol of a primary alcohol (either ethanol or butanol) requires 2 mol of NADH, and each glucose consumed produces 2 mol of NADH, *at least one mol of primary alcohols must be produced per mol glucose consumed to satisfy the electron balance when NADH cannot be converted to H_2_* (Table 2). This “minimum primary alcohol yield” applies to 4-carbon knockout strains, such as *C. acetobutylicum* strain CACas9 *Δhbd* p95ace02_atoB, when they are grown on glucose with exogenous butyrate.

As expected, the *C. acetobutylicum* monoculture grown under a N_2_ headspace produced large quantities of H_2_ and CO_2_ (Fig. 2C). The primary alcohol yield of this culture was 1.07 mol ethanol per mol glucose (1.02 mol ethanol per mol glucose) (Fig. 2B), close to the minimum primary alcohol yield of 1 mol primary alcohol per mol glucose for this strain expected under the assumption of no NADH to H_2_ conversion (Table 2). In the corresponding coculture control grown under a N_2_ headspace, CO_2_ was produced, but there was no detectable H_2_ accumulation (Fig. 2D). This is a significant result because it shows that, in the coculture, all the electrons from glucose are captured as soluble products; no electrons are converted to gaseous H_2_, which is viewed as a waste product and a longstanding drawback of classical ABE fermentation. Strikingly, this coculture produced a primary alcohol yield of 0.83 mol per mol glucose (0.78 ethanol, 0.05 butanol) (Fig. 2B), well below the minimum yield of 1 mol primary alcohol per mol glucose (Table 2). This result requires interspecies electron transfer. To demonstrate that, we carried out electron balances based on the established metabolic stoichiometry (as summarized in Table 2) and the analysis above, as follows.

We consider the glycolytic stoichiometry (Eq. T1, Table 2), the stoichiometric reactions for *C. acetobutylicum* production of ethanol from acetyl-CoA (Eq. T4, Table 2) and butanol from butyrate (Eq. T5, Table 2). Using the per-glucose ethanol and butanol yields (0.78 and 0.05, respectively) from the coculture under a N_2_ headspace (Fig. 2B), we write Eqs. 3 and 4,

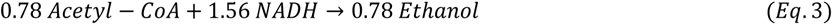

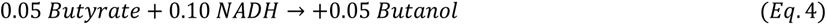

and combining with the glycolytic stoichiometry, we then write Eq. 5,

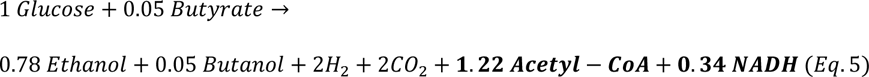

which shows that, after subtracting the acetyl-CoA and NADH used for ethanol and butanol production, 1.22 mol acetyl-CoA and 0.34 mol NADH per mol glucose (17% of the NADH generated from glycolysis) were used in other biological processes.

Acetyl-CoA is used for acetate or acetone production (Table 2, Eqs. T8, T9). Per the analysis above, since monocultures of the *C. acetobutylicum* strain CACas9 *Δhbd* p95ace02_atoB cannot produce H_2_ using NADH, this *C. acetobutylicum* strain must dispose of the NADH in coculture via a different mechanism. Based on the carbon balance (using glucose consumption and optical density measurements), only 3% of metabolized glucose was utilized for biomass formation (as is consistently observed for *C. acetobutylicum* fermentations [3]), so only 0.03 mol per mol glucose of the unaccounted for NADH can be explained via biomass formation (Table 2, Eq. T3). This leaves 0.31 mol NADH per mol glucose (16% of NADH generated from glycolysis) unaccounted for.

Based on these NADH balances we can conclude that coculture with *C. ljungdahlii* enables *C. acetobutylicum* to either convert NADH into H_2_ (which will be assimilated by *C. ljungdahlii* once released into the medium), pass electron carriers directly to *C. ljungdahlii* via cytoplasmic exchange, or both, as these two mechanisms are not mutually exclusive. The absence of detectable gaseous H_2_ in the coculture under a nitrogen atmosphere (Fig. 2) suggests that direct electron transfer may be involved.

The cocultures grown under H_2_ and the H_2_/CO_2_ mix show significant net consumption of H_2_ (Fig. 2E, F), suggesting that the cultures are electron limited and capable of assimilating electrons above and beyond what is available from glucose catabolism. This is a crucial finding because, in traditional sugar fermentations, the degree of reduction of all metabolic products taken together is limited by the available electrons in the sugar substrate [22]. Here, we demonstrate that this requirement can be overcome in coculture by supplying additional reducing equivalents in the form of H_2_.

Significantly, the addition of H_2_ and CO_2_ to the headspace led to an increase in isopropanol yield relative to the nitrogen and H_2_ headspace controls (0.77 mM isopropanol per mol glucose compared to 0.56 and 0.61, respectively) (Fig. 2B). This condition also resulted in the lowest primary alcohol yield (a desirable feature from the point of view of isopropanol production) of the three coculture headspace conditions tested (0.71 mM primary alcohols per mM glucose) (Fig. 2B). Since the addition of both H_2_ and CO_2_ enables *C. ljungdahlii* to initiate growth via its autotrophic metabolism immediately from the beginning of the fermentation (rather than after *C. acetobutylicum* starts generating CO_2_), these results suggest that maximizing the availability of gaseous substrates for *C. ljungdahlii* helps to minimize the ethanol yield and maximize the isopropanol yield from the coculture.

### 3.4. “Pseudo-perfusion” experiments demonstrate that high-density cocultures with supplemental *C. ljungdahlii* increase interspecies electron transfer

Next, we tested the impact of higher cell densities on the electron management/transport discussed above and the associated metabolic behavior of the coculture. We hypothesized that higher cell densities would enable “tighter” cell-to-cell interactions and more efficient electron exchange, leading to higher isopropanol yields and decreased ethanol production. To test this hypothesis, we employed an established “pseudo-perfusion” protocol [23] using cell concentration and resuspension, as follows. We inoculated a coculture of CACas9 Δ*hbd* p95ace02_atoB and *C. ljungdahlii* p100ptaHalo at a relatively high starting cell density of approximately OD_600_ = 6, with a 5:1 ratio of *C. ljungdahlii* to *C. acetobutylicum*, in 20 mL of TCGMB growth medium with 80 g/L glucose and 1 g/L fructose in 160 mL serum bottles. Then, approximately every 8-12 hours, we pelleted the cells, decanted the spent media, and resuspended the cells in fresh medium. This approach enabled us to quickly and easily simulate the high cell densities characteristics of high-density perfusion cultures.

In addition, at every resuspension, we flushed and pressurized the serum bottle headspace with 20 psig of 80/20 H_2_/CO_2_ gas mix thus providing extra gaseous electron and carbon substrates to support *C. ljungdahlii’*s growth and reduction of acetone to isopropanol. We also added more *C. ljungdahlii* cells every 8-12 hours with every fresh-medium resuspension in order to maintain a good ratio of the two species population [3], given that *C. ljungdahlii* grows much slower than *C. acetobutylicum*. In one experimental set, we spun down and added 20 mL of ∼1 OD_600_ fresh *C. ljungdahlii* cells at each resuspension and, in the second set, we spun down and added twice as much, 40 mL of ∼1 OD_600_ fresh *C. ljungdahlii* cells. The pseudo-perfusion strategy is illustrated schematically in Figure S3.

When interrogating metabolite yields from the pseudo-perfusion experiments (and subsequent perfusion experiments), we report, in addition to the isopropanol and primary alcohol yields, the combined yield of isopropanol and acetone as the “3C” metabolite yield to more concisely and accurately communicate the total molar amount of acetone produced per mol glucose by *C. acetobutylicum,* as WT *C. ljungdahlii* cannot synthesize acetone. In coculture, most acetone is reduced to isopropanol by *C. ljungdahlii*.

In both sets of pseudo-perfusion conditions, we observed very high biomass accumulation of OD_600_ >40 (Fig. 3), which is approximately 4-fold higher than the cell densities of 8-12 OD_600_ typically achieved in batch coculture experiments (Fig. 3A, C). Adding *C. ljungdahlii* cells at an OD_600_ of ∼2 resulted in strong isopropanol and acetone yields (up to 0.98 mol 3C products per mol glucose) and corresponding decreases in primary alcohol yield (as low as 0.16 mol primary alcohols per mol glucose), especially across the last four resuspension periods (Fig. 3D). Adding *C. ljungdahlii* cells at an OD_600_ of ∼1 resulted also in decreasing primary alcohol yields and increasing 3C product yields with each successive resuspension (Fig. 3B), but the results were more modest. These findings suggest that building the cell density through the pseudo-perfusion strategy, combined with the addition of extra *C. ljungdahlii*, could redirect up to 84% of the excess NADH from glycolysis (based on the lowest demonstrated primary alcohol yield) from *C. acetobutylicum* to *C. ljungdahlii*, either via NADH to H_2_ conversion, direct electron carrier transfer, or both.

**Figure 3:**
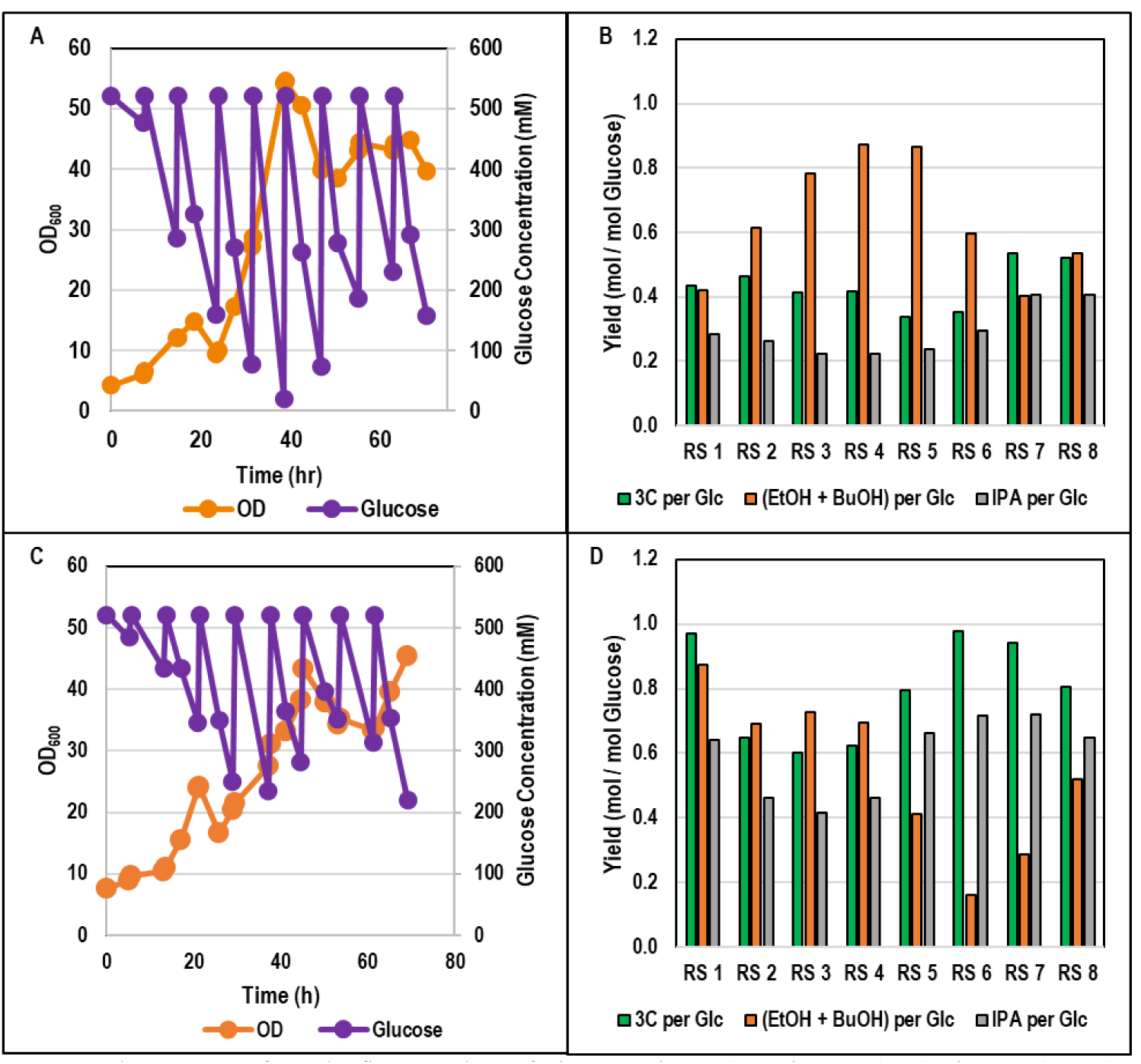
The outcomes from the first pseudo-perfusion experiment (see Figure S3). A) Biomass (OD600) and glucose profiles for 1 OD600 of fresh *C. ljungdahlii* cells added at each resuspension. B) Major metabolite yields for the experiment of panel A. C) Biomass (OD600) and glucose profiles for 2 OD600 of fresh *C. ljungdahlii* cells added at each resuspension. D) Major metabolite yields for the experiment of panel C.

### 3.5. Increased *C. ljungdahlii* population fractions give rise to high 3C yields and sugar-carbon recoveries into metabolites

To further explore these findings, we performed additional pseudo-perfusion cocultures with 40 mL of ∼2 OD_600_ fresh *C. ljungdahlii* added at every resuspension. We consistently observed the same trend of decreasing primary alcohol yields with continuing resuspension rounds. We also observed increasing 3C yields from the later resuspension periods, with maximum 3C yields of 1.39 and 1.57 mol per mol glucose across the last two resuspension periods, respectively, for a period of 19 hours (Fig. 4A), the highest 3C yields we had ever observed from a *C. acetobutylicum-C. ljungdahlii* coculture up to that point. To further interrogate this finding, we quantified the population distribution throughout the course of this pseudo-perfusion experiment using RNA-FISH [2]. These data show that the highest selectivity for 3C metabolites (and lowest primary alcohol yields) correspond to the timepoints with the highest *C. ljungdahlii* cell fraction (Fig. 4C). Notably, across these last two resuspension periods (19 hours of growth) with high *C. ljungdahlii* cell fractions, we also achieved carbon-neutral or carbon-negative glucose fermentation, re-assimilating all CO_2_ from glycolysis, along with additional exogenous CO_2_ from the headspace, into metabolic products (Fig. 4D). These findings support our hypothesis that strong autotrophic activity of *C. ljungdahlii* results in higher 3C and isopropanol yields from *C. acetobutylicum*.

**Figure 4:**
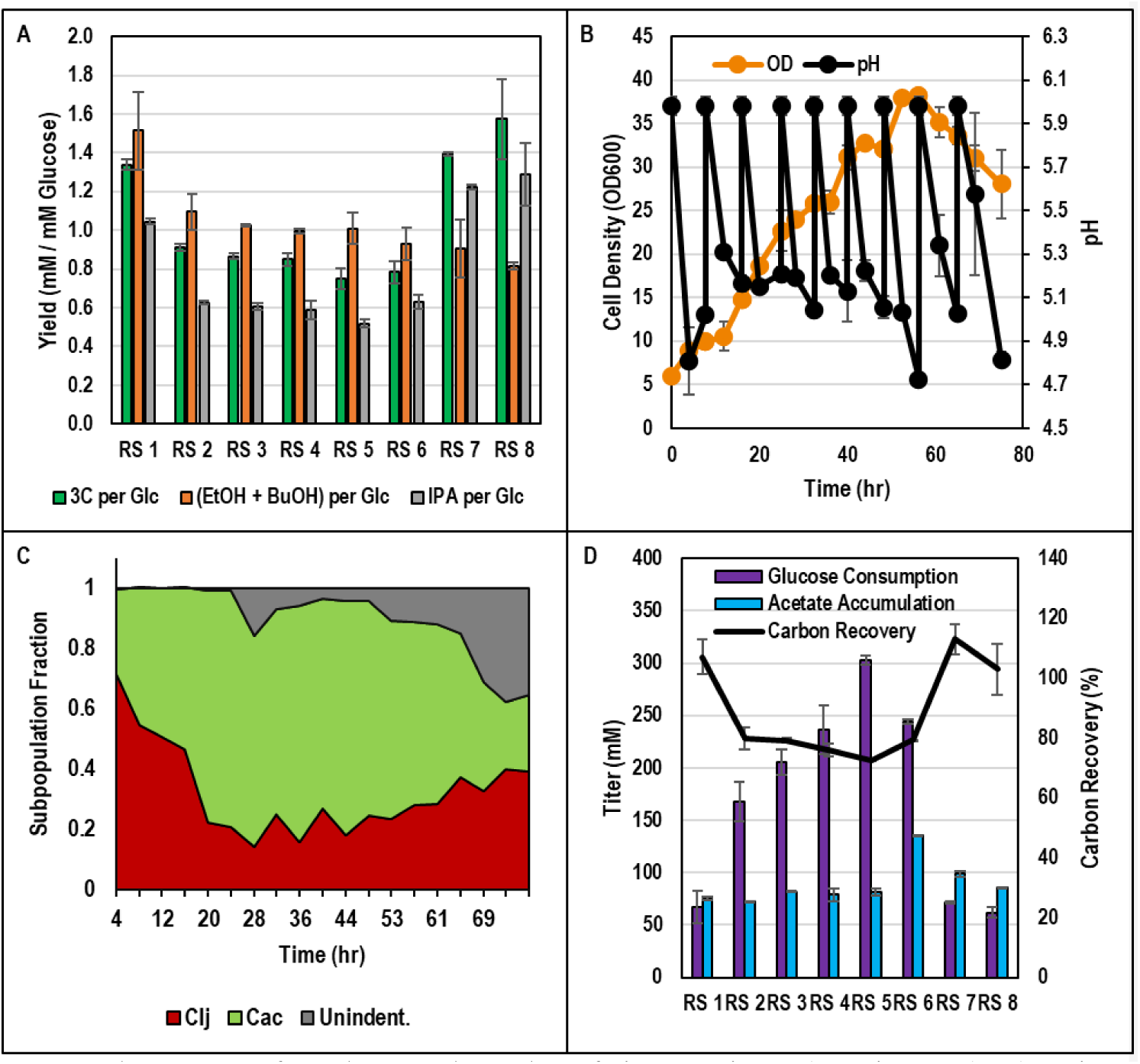
The outcomes from the second pseudo-perfusion experiment (see Figure S3). A) Major metabolite yields. B) Biomass (OD600) and pH profiles. C) *C. acetobutylicum* and *C. ljungdahlii* population kinetics. D) Glucose consumption, acetate accumulation, and carbon recovery. Error bars represent standard deviations from two biological replicates.

Despite the high selectivity for isopropanol, we observed decreased glucose utilization across the last two resuspensions (Fig. 4D). Based on the established pH-dependent inhibitory effect of acetate on *C. acetobutylicum* growth [24–26], inhibition of glucose consumption is likely due to the accumulation of high levels of acetate in the coculture, due to robust CO_2_ fixation by *C. ljungdahlii,* combined with the high pH variability of our pseudo-perfusion experiments (Figs. 3A, C & Fig. 4B). We supported this assumption by additional data which showed that high levels of exogenous acetate fed to *C. acetobutylicum* monocultures inhibit biomass accumulation and glucose consumption (Fig. S4).

### 3.6. Coculture in a high density pH-controlled perfusion reactor enabled carbon-negative glucose fermentation with isopropanol as the sole alcohol produced from glucose

Based on the pseudo-perfusion experiments, we concluded that increasing the coculture cell density, combined with accumulating large fractions of active *C. ljungdahlii* cells, would enable high isopropanol yields from glucose. However, accumulation of toxic levels of acetate prevented complete consumption of glucose by *C. acetobutylicum* during the pseudo-perfusion periods in which we observed the highest isopropanol yields. To overcome this problem, we used a pH-controlled bioreactor operated in continuous mode with cell retention (perfusion operation). An in-line packed bed column was integrated specifically to support strong autotrophic growth and activity of *C. ljungdahlii* via efficient gassing of external H_2_ and CO_2_ (Fig. 5A). The feed concentration of glucose (110 mM) and the reactor dilution rate (0.083 hr^-1^; 2 vessel volumes per day) were chosen to achieve strong isopropanol productivity while minimizing accumulation of acetate to avoid inhibiting *C. acetobutylicum* (Supplementary Text Document 1).

**Figure 5:**
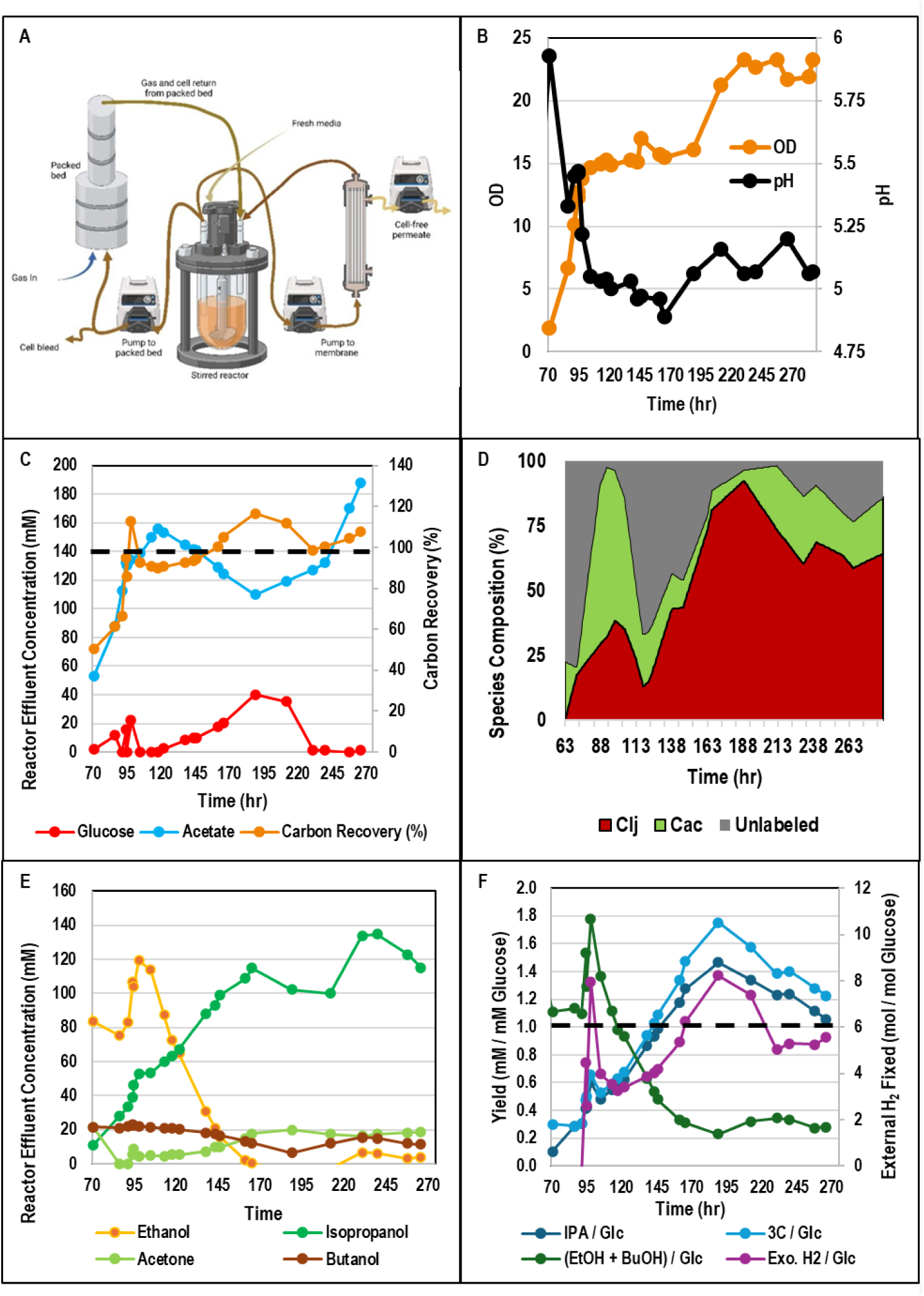
The first perfusion experiment. A) Schematic of perfusion bioreactor configuration. B) Biomass (OD600) and pH kinetics. C) Glucose, acetate, and carbon recovery kinetics. D) *C. acetobutylicum* and *C. ljungdahlii* population kinetics. E) Isopropanol, acetone, ethanol, and butanol kinetics. F) Yields of 3C metabolites, isopropanol, and primary alcohols and external H2 fixed per mol glucose.

*C. acetobutylicum* (CACas9 Δ*hbd* p95ace02_atoB) was inoculated first to reduce glucose concentration before the addition of *C. ljungdahlii* (carrying the p100ptaHALO plasmid for the antibiotic resistance) approximately 63 hours into the run. Residual glucose concentrations dropped to virtually zero around hour 100 before starting to rise again after 170 hours and reaching levels between 20 and 40 mM (indicating 64-82% glucose utilization, based on the 110 mM glucose in the feed) between ca. 170 and 220 hours (Fig. 5C). This period of decreased glucose utilization closely correlated with a very low fraction of *C. acetobutylicum* cells in the co-culture population (Fig. 5D), likely caused by the acetate accumulation, peaking at 118 hours at 156 mM, a toxic level at the corresponding pH of 5.04 (Fig. 5B, C). Starting at 189 hours, the *C. acetobutylicum* population began to recover. More than 98% of available glucose was utilized for the final 35 hours of the experiment (Fig. 5C).

Using an electron or “degree of reduction” balance [22] these data show that in addition to the H_2_ produced from glucose catabolism by *C. acetobutylicum,* the coculture fixed up to 8 mol of external H_2_ per mol glucose consumed, meaning that up to 40% of electrons in the products originated from external H_2_ (Fig. 5F). This represents a powerful demonstration of the ability of the *C. acetobutylicum-C. ljungdahlii* coculture to co-utilize additional reducing equivalents (electron) sources alongside glucose.

At approximately 100 hours, following the initial biomass accumulation phase, ethanol production started to decrease rapidly. After about 50 hours, the primary alcohol yield stabilized between 0.2-0.4 mol per mol glucose, with 10 mM or less of accumulated ethanol, for the rest of the fermentation (Fig. 5E, F). These primary alcohol yields require that, for the final 104 hours of fermentation, 60-80% of NADH from glycolysis was passed from *C. acetobutylicum* to *C. ljungdahlii* via either extracellular H_2_ or intercellular direct exchange (or both).

The robust interspecies exchange of electrons enabled very high isopropanol and 3C yields. As the ethanol concentration began to decrease, the isopropanol concentration sharply rose, climbing to 115 mM by 166 hours and eventually peaking at 135 mM (representing a volumetric productivity of 11.2 mM isopropanol per hour) at 240 hours (Fig. 5E). The maximum observed yields of isopropanol (1.47 mol per mol glucose) and 3C metabolites (1.75 mol per mol glucose) were observed at 189 hours (Fig. 5F), higher than any of the yields previously achieved from batch or pseudo-perfusion fermentations. For comparison, this 3C yield is 2.9-fold and 3.5-fold the maximum acetone yields per glucose previously determined for *C. acetobutylicum* monoculture strains with (0.6 mol acetone per mol glucose) and without 4-carbon metabolism (0.5 mol acetone per mol glucose), respectively (Eqs. 1 & 2).

Yields of total 3C metabolites remained at or above 1.2 mol per mol glucose for the final 100 hours of the fermentation (Fig. 5F), indicating strong acetate uptake by *C. acetobutylicum* as explained next. The maximum yield of acetone from glucose by *C. acetobutylicum* (assuming the excess NADH generated by glycolysis is removed by *C. ljungdahlii*) is 1.0 mol per mol glucose [22] based on the fact that a very small fraction of glucose is used for biomass formation. Therefore, 3C yields over this value (as observed in this fermentation) requires uptake of acetate produced from CO_2_ fixation by *C. ljungdahlii*. Our maximum observed yield of 1.75 mol 3C metabolites per mol glucose (Fig. 5F) indicates that as much as 1.5 mol acetate was assimilated (to produce acetone) per mol glucose consumed.

Although acetate concentrations were maintained at levels low enough to allow robust *C. acetobutylicum* activity throughout the fermentation, accumulation of acetate climbed to 188 mM (376 C-mM) late in the experiment, making it the second major product behind isopropanol (405 C-mM) on a carbon mole basis (total 3C metabolites peaked at 459 C-mM) (Fig. 5C, E). This was largely due to the exceptionally high 3C yields (and concomitant production of CO_2_ which accompanies acetone production due to the decarboxylation of acetoacetate, Fig. 1A) and the strong fixation of external CO_2_ by *C. ljungdahlii*, clearly illustrated by the exceptional carbon recovery of the fermentation. **For the final 104 hours, we demonstrated carbon-negative glucose fermentation (incorporation of all glucose carbon and additional exogenous CO_2_ into metabolites), with a maximum demonstrated carbon recovery of 117%** (at 189 hours), at all but one timepoint (99% carbon recovery at 231 hours) (Fig. 5C).

This period of carbon-negative fermentation corresponded with the time points during which 50% or more of cells in the reactor were *C. ljungdahlii* (Fig. 5D). This timeframe also included the peak isopropanol and 3C yields, further building upon previous batch and pseudo-perfusion data suggesting that strong autotrophic activity by *C. ljungdahlii* (as indicated by the carbon recovery) and high cell fraction of *C. ljungdahlii* (as indicated by the population data) enable exceptional isopropanol and 3C yields from the coculture. Notably, for two consecutive timepoints within this *C. ljungdahlii* dominant period (representing 23 hours of fermentation), production of ethanol was eliminated (Fig. 5E); isopropanol was the sole alcohol produced from glucose during this time. This phenotype was due to the robust transfer of electrons from *C. acetobutylicum* to *C. ljungdahlii* (via extracellular H_2_, intercellular direct exchange, or both) enabled by the high cell density conditions achieved and stably maintained in the perfusion reactor.

## CONCLUSIONS

Genetically modifying single organisms for simultaneous utilization of diverse substrates, while maintaining high yield and productivity of a target product, can be extremely challenging due to both methodological reasons (especially for non-model bacteria) and fundamental stoichiometric and thermodynamic pathway limitations. Here, we demonstrate that implementing a coculture strategy between *C. acetobutylicum* and *C. ljungdahlii* in a high-density perfusion fermentation enables acetone yields from an engineered *C. acetobutylicum* strain (converted to isopropanol in the coculture) that cannot be achieved in monocultures of the same strain (or even using the same coculture approach in batch fermentations at lower cell density). Based on the well-established stoichiometry of the key metabolic reactions of both *C. acetobutylicum* and *C. ljungdahlii*, we show that the isopropanol yields demonstrated here require enhanced electron exchange between the two organisms via extracellular H_2_ and/or intercellular electron carriers. Finally, we were able to simultaneously demonstrate robust assimilation of CO_2_ and H_2_ by *C. ljungdahlii* in coculture, achieving “carbon-negative” glucose fermentation in which all carbon and electrons in the glucose substrate are assimilated to products alongside large amounts of external CO_2_ and H_2_.

In addition to the perfusion fermentation run discussed in detail in section 3.6, we have consistently shown that prolonged continuous *C. acetobutylicum-C. ljungdahlii* coculture fermentations with cell retention lead to production of isopropanol as the major or sole alcohol product from the coculture (one such example, which tested progressive increase in glucose feed rates, is presented in Figure S5). The major rate-limiting factors for scaling of this system to produce economically viable isopropanol productivities are the ability of *C. acetobutylicum* to assimilate more of the acetate produced by *C. ljungdahlii*, which is consistently the second major product behind isopropanol, and to improve the gas transfer of the bioreactor system to enable sufficient transport of CO_2_ and H_2_ to enable carbon-negative fermentation at industrially relevant glucose feed rates. For acetate uptake, we suspect that the activity of the native *C. acetobutylicum* coenzyme-A transferase (ctfA and ctfB), which has a notably high K_M_ for acetate (1200 mM) [27], is likely limiting; future optimization of the coculture will evaluate enzymes with higher activity towards acetate. With respect to improving gas transfer, scaling and optimizing industrial gas fermentation is an active area of interest in both the academic and industrial research communities [28].

The syntrophic coculture between *C. acetobutylicum* and *C. ljungdahlii* represents a promising approach to enable 1) the renewable production of chemicals with a degree of reduction greater than glucose via assimilation of additional reducing equivalents and 2) the co-utilization of external CO_2_ alongside biomass sugars, contributing to a sustainable, carbon-negative bioeconomy. The unique properties of this system, specifically enhanced electron exchange at higher cell densities, are also worthy of further study, and may have important implications for the implementation of cocultures in industrial biotechnology as well as improved understanding of electron exchange mechanisms in both natural and synthetic microbial communities.

## SUPPLEMENTAL MATERIAL

**Supplemental Figures (Word document)** (Figures S1-S5)

**Supplementary Text Document 1 (Word document)** (“Estimating Acetate Accumulation for Experimental Design of Coculture Perfusion Experiment”)

## ACKNOWLEDGEMENTS

This work was supported by an ARPA-E project under contract AR0001505. N.B.W. and J.D.H. were supported in part by a U.S. Department of Education GAANN Fellowship under grant P200A210065.

## Author contributions

N.B.W., J.K.O., H.S., and E.T.P. designed research; N.B.W., J.K.O., H.S., P.C.M., and J.D.H. performed research; N.B.W., J.K.O., J.D.H., and E.T.P. analyzed data; E.T.P. supervised the project; and N.B.W., J.K.O. and E.T.P. wrote the paper.

---

**Table S1.**
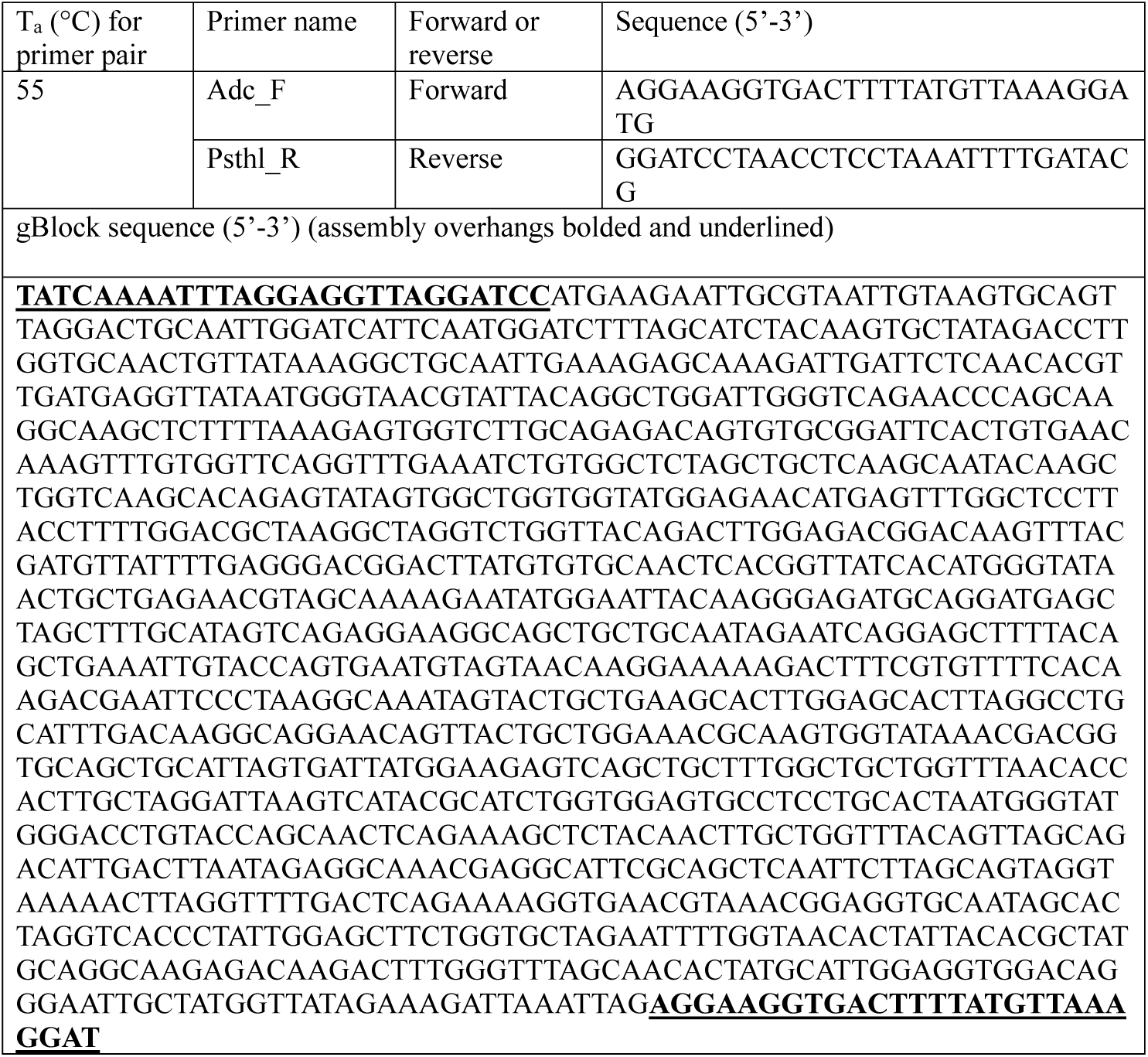
Primers used for cloning.

**Figure S1:**
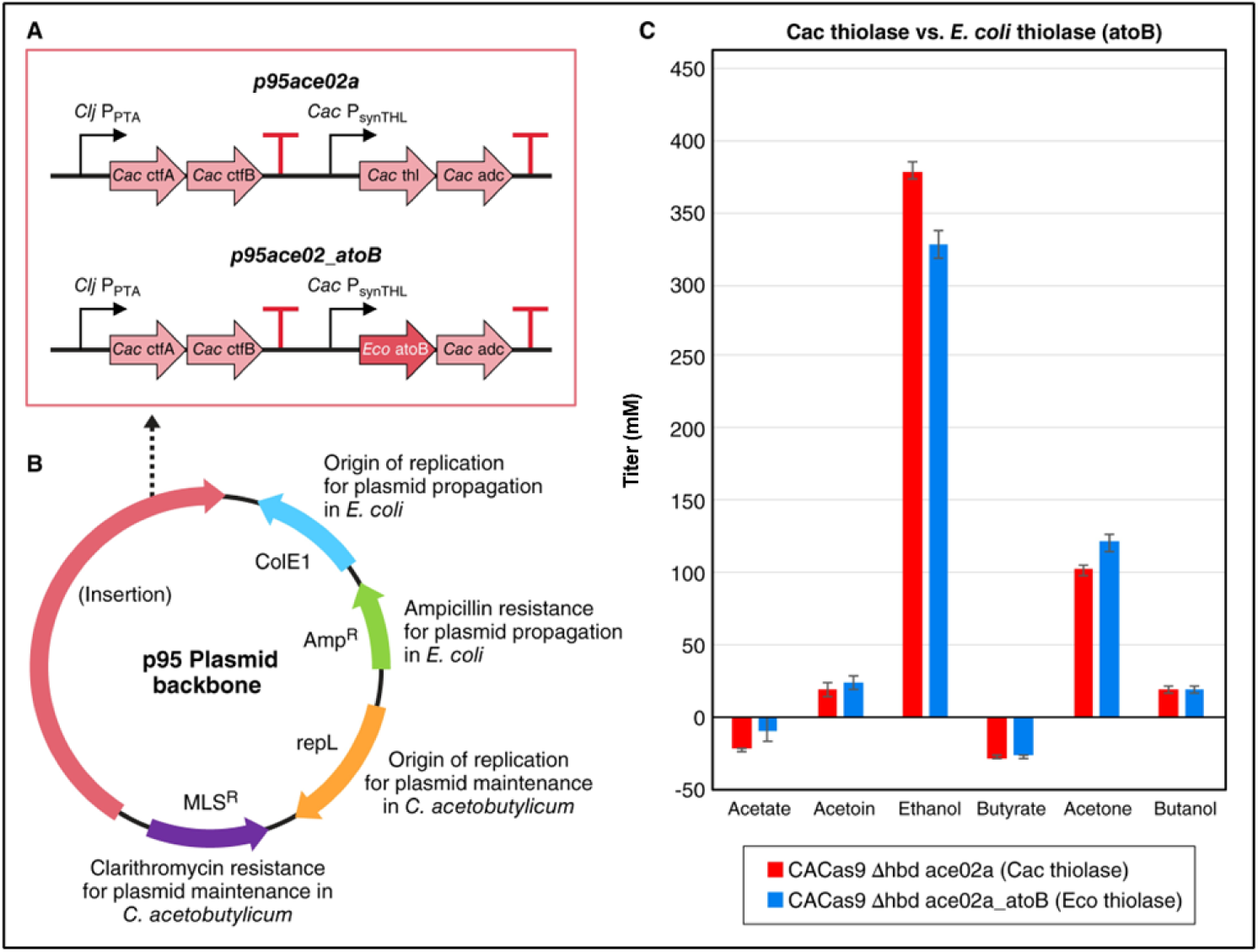
A) Schematic comparing acetone operon construction between p95ace02a and p95ace02_atoB acetone plasmid variants. B) Schematic of p95 *C. acetobutylicum* shuttle vector organization. C) Comparison of major metabolite titers between monocultures of *C. acetobutylicum* (CACas9 Δ*hbd*) carrying either p95ace02a or p95ace02_atoB.

**Figure S2:**
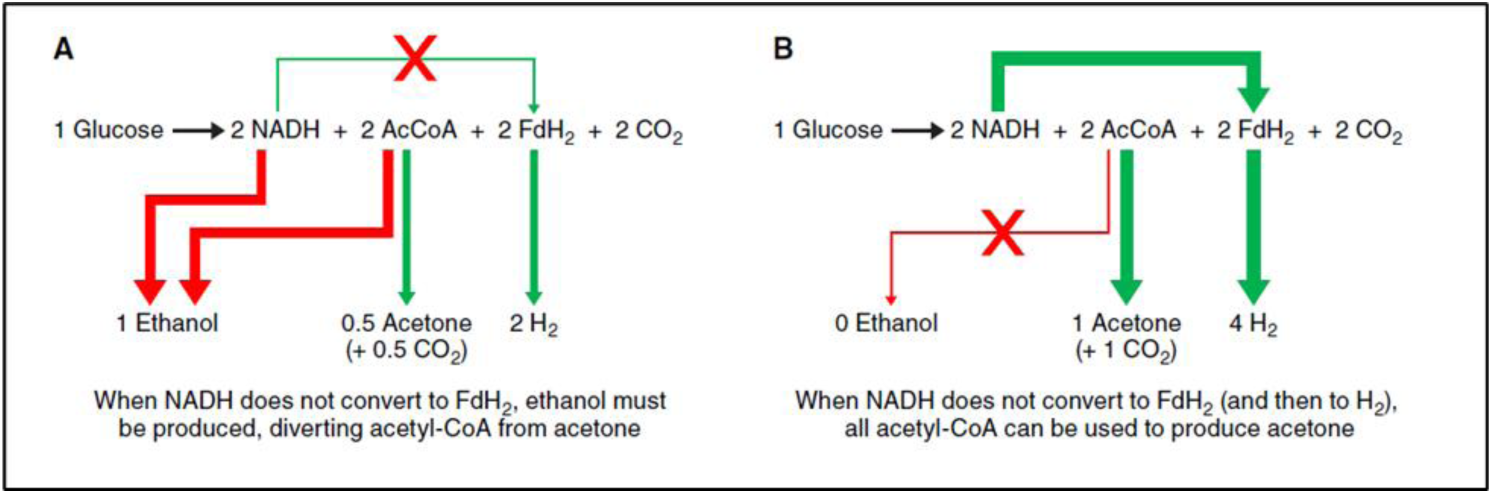
A) Schematic of electron and carbon flows when NADH cannot be converted to ferredoxin. B) Schematic of electron and carbon flows when NADH can be converted to ferredoxin and then to H2.

**Figure S3:**
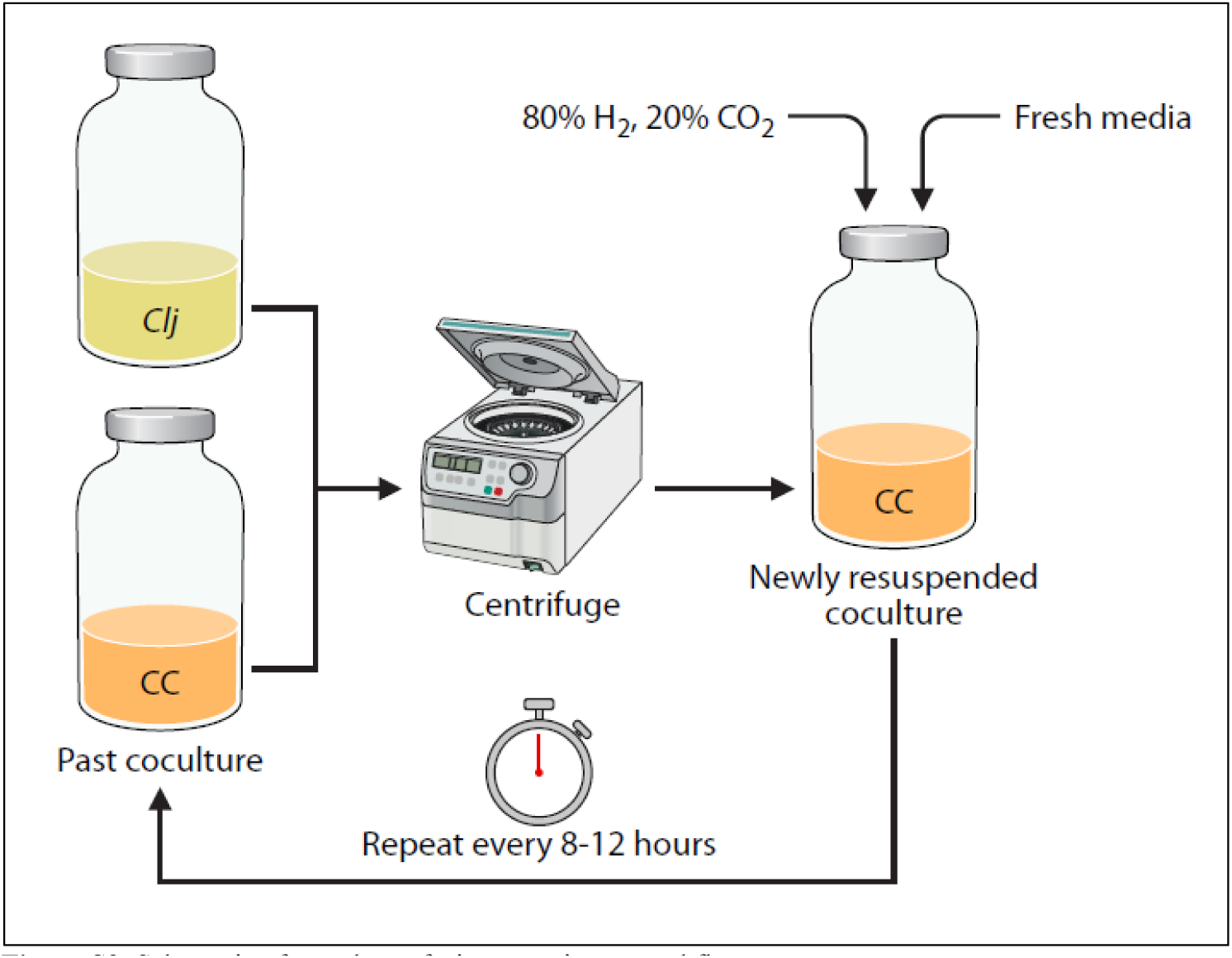
Schematic of pseudo-perfusion experiment workflow.

**Figure S4:**
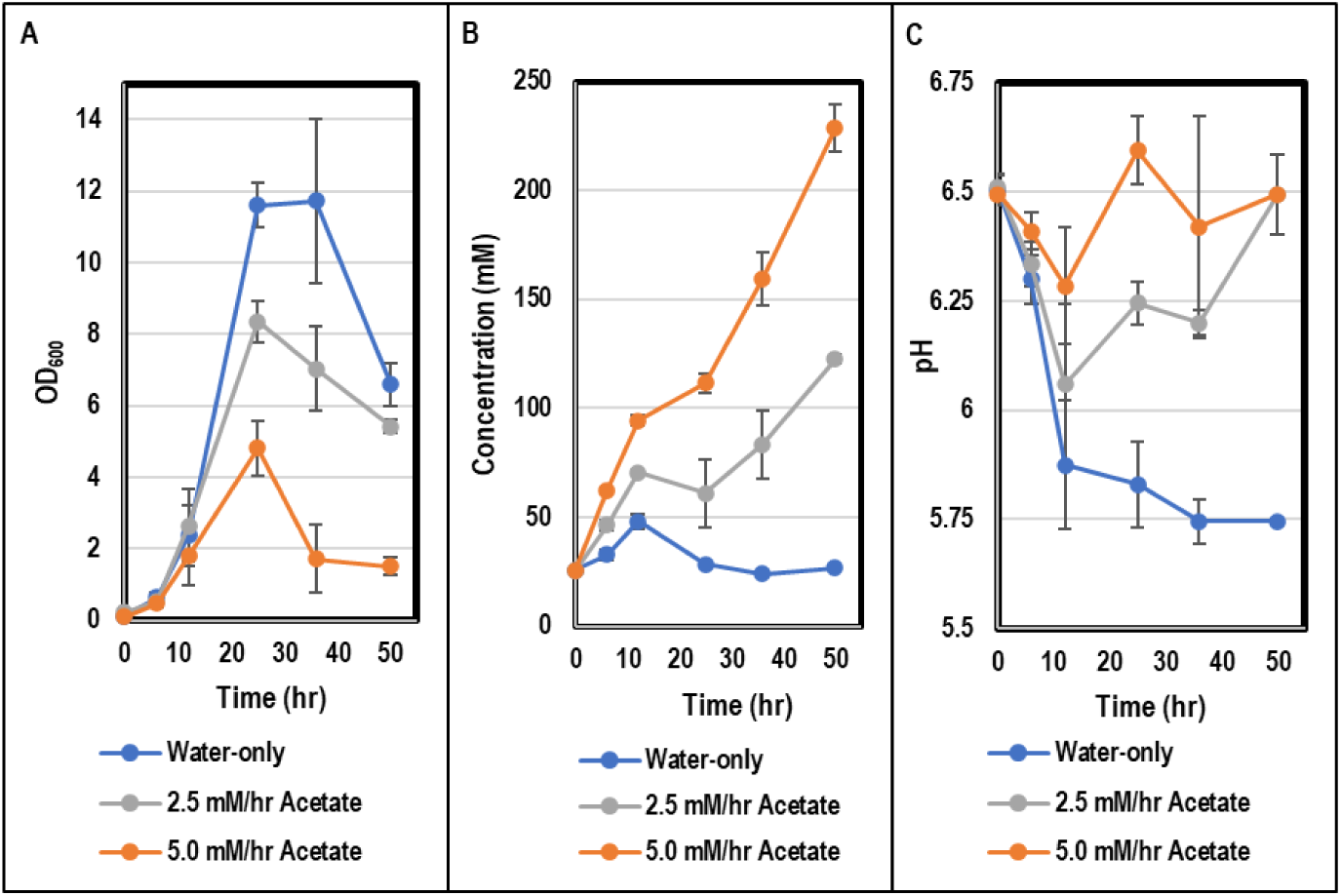
A) Biomass (OD600) kinetics of *C. acetobutylicum* (CACas9 Δ*hbd* p95ace02_atoB) monocultures fed water, 2.5 mM acetate per hour, or 5.0 mM acetate per hour. B) Acetate kinetics in the water, 2.5 mM acetate per hour, and 5.0 mM acetate per hour conditions. C) pH kinetics in the water, 2.5 mM acetate per hour, and 5.0 mM acetate per hour conditions.

**Figure S5:**
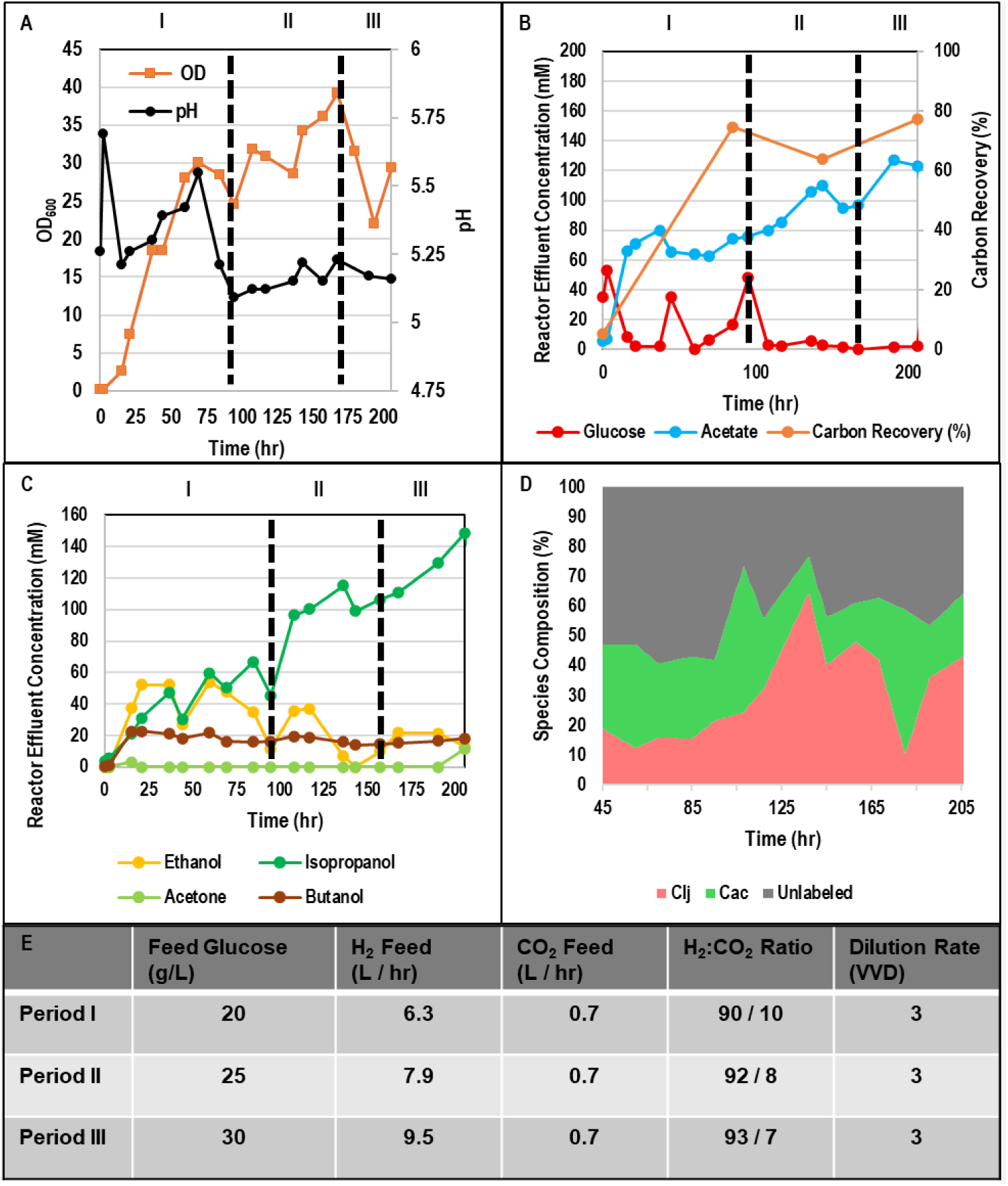
The second perfusion experiment. A) Biomass (OD600) and pH kinetics. B) Glucose, acetate, and carbon recovery kinetics. C) Isopropanol, acetone, ethanol, and butanol kinetics. D) *C. acetobutylicum* and *C. ljungdahlii* population kinetics. E) Glucose feed concentrations, gas flow rates, and reactor dilution rates for fermentation periods I-III.

## Notes

### Competing Interest Statement

The authors have declared no competing interest.

