## Supplemental Text Document 1 for "Enabling supratheoretical isopropanol yields from carbon-negative glucose fermentations with *Clostridium acetobutylicum*-*Clostridium ljungdahlii* cocultures"

**Supplementary Document 1**

**Estimating Acetate Accumulation for Experimental Design of Coculture Perfusion Experiment**

In the pseudo-perfusion experiments we started to observe decreases in the rate of glucose utilization by *C. acetobutylicum* when acetate accumulated in excess of 100 mM. Since pyruvate decarboxylation produces 2 CO_2_ per glucose, and 2 CO_2_ are combined to produce 1 acetate by *C. ljungdahlii*, we estimated that a glucose feed concentration of 110 mM (20 g/L) would maintain the in-situ acetate concentration at approximately 110 mM (assuming continuous fermentation where *C. ljungdahlii* reduces most or all of the excess CO_2_ from glycolysis into acetate).

This value was only intended to be a ballpark estimate of acetate accumulation, not a precise carbon balance. For example, we could not account for the extra CO_2_ generated by *C. acetobutylicum* from acetoacetate decarboxylation (in the acetone pathway) because we could not know *a priori* the acetone yield per glucose. Similarly, we could not predict how much, if any, exogenous CO_2_ *C. ljungdahlii* would assimilate from the external gas feed (via the packed bed column). Both of these additional CO_2_ sources had the potential to increase acetate accumulation above 110mM. Conversely, *C. acetobutylicum* has the capacity to assimilate acetate into the acetone pathway, and it was possible that *C. ljungdahlii* would not completely assimilate the excess CO_2_ from glycolysis. Both of these factors would exert downwards pressure on acetate accumulation. Finally, we also hypothesized that *C. acetobutylicum* acetate tolerance would improve at the controlled, stable pH (setpoint of 5.1) of the perfusion bioreactor system relative to *C. acetobutylicum* acetate tolerance under the highly variable pH conditions of the pseudo-perfusion experiments. Based on these considerations, 110 mM (20 g/L) of glucose was chosen as an input feed concentration which would likely prevent acetate from accumulating to levels inhibitory to *C. acetobutylicum* under the conditions tested.

For typical batch experiments, we provide the coculture with 80 g/L of glucose (440 mM) in the medium, most or all of which is consumed over the course of 36-72 hours (depending on the strain and growth conditions). Consumption of 440 mM of glucose in 48 hours corresponds to an average glucose consumption rate of 9.17 mM glucose per hour. To demonstrate a similar rate of consumption in the perfusion reactor, we set the reactor dilution rate to 0.083 hr^-1^ (2 vessel volumes per day) which, when combined with the input feed concentration of 110 mM glucose, delivered a volumetric glucose feed rate of 9.13 mM glucose per hour to the reactor.
